## Supplementary figures and images for "Interplay of human macrophage response and natural resistance of *L.* (*V*.) *panamensis* to pentavalent antimony"

### Figure 1S

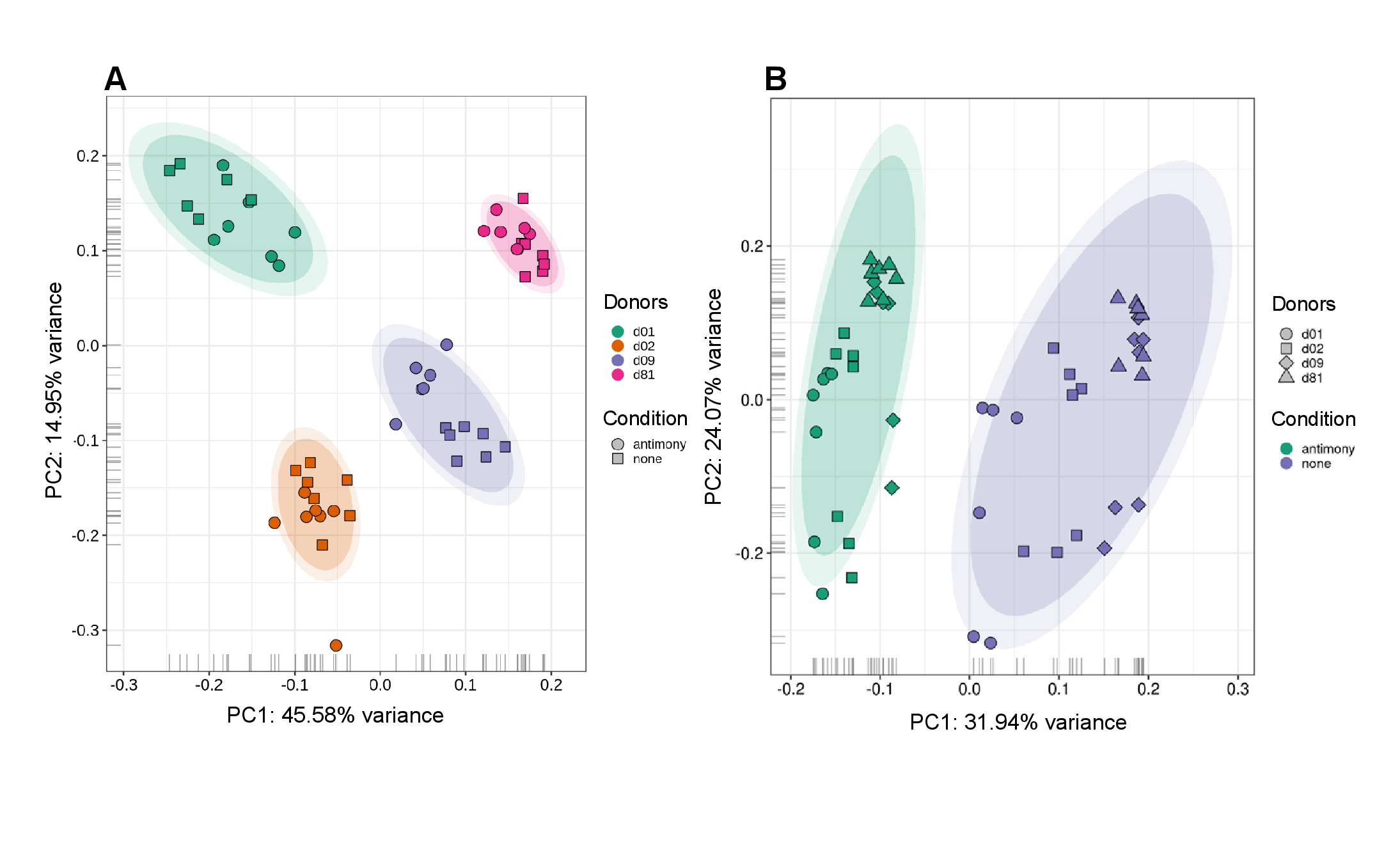

### Figure 2S

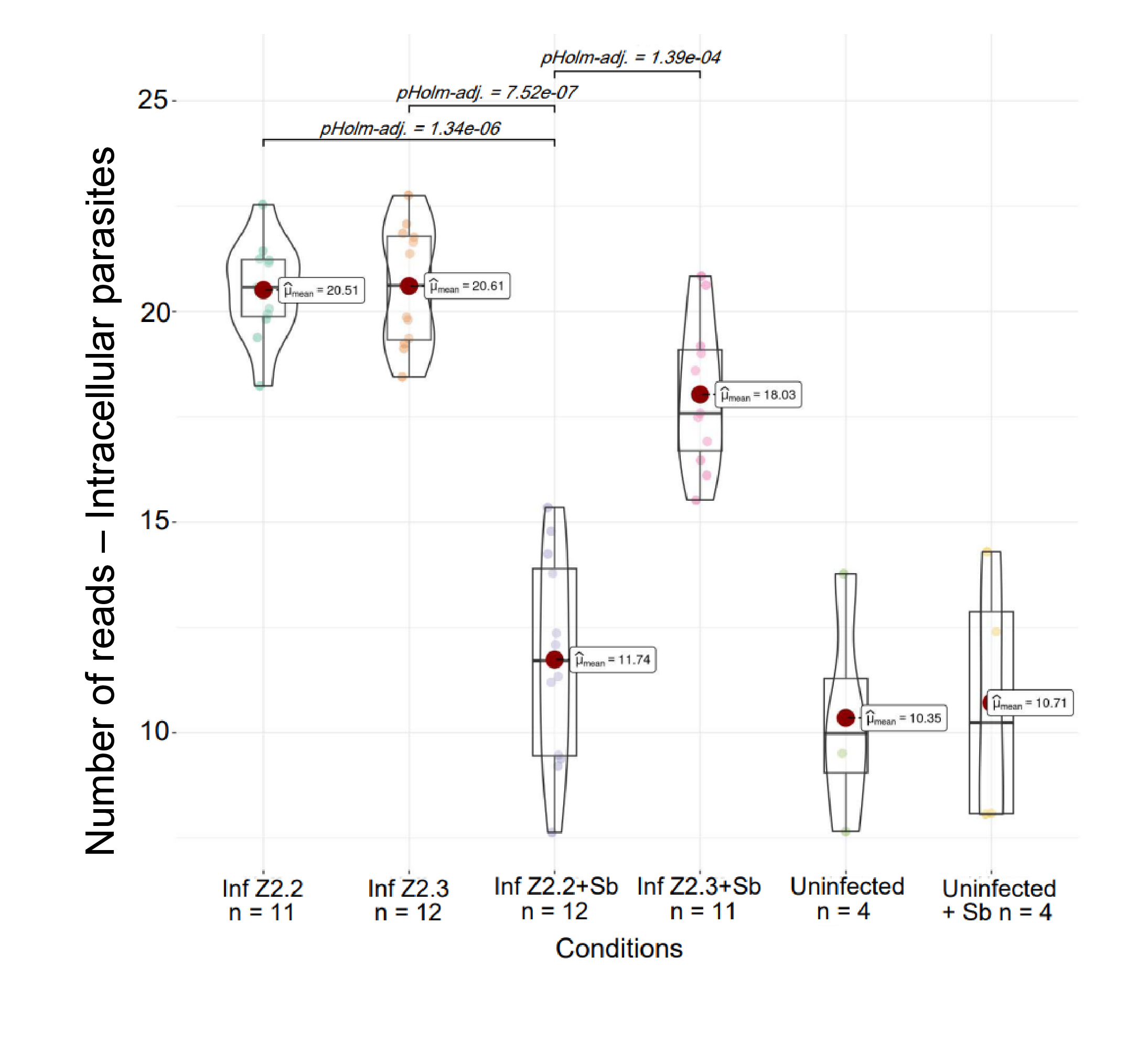
